# Characteristic impairments of goal-directed and habitual control in bulimia nervosa

**DOI:** 10.1101/348250

**Authors:** Abraham Nunes, Thomas J. Helson, Laura M. Dixon, Thomas P. Trappenberg, Aaron R. Keshen

## Abstract

The relationship between clinical eating disorder symptom severity and balance of model-based (MB) and model-free (MF) control is unclear, and these traits’ predictive capacity is untested in this population. In 25 healthy controls (HCs), 25 subjects with binge eating disorder (BED), and 25 subjects with bulimia nervosa (BN), we show an inverse relationship between symptom severity and MB (though not MF) control. However, trial-by-trial behavioural data discriminated BN from other groups (area under receiver operating characteristic curve of 0.78; 95% CI=0.64–0.92) based primarily on impaired MF control. Our data—including analyses of reaction time and theory-driven computational modeling—support the hypothesis that among pathological binge eating groups, BN may be characterized by impaired value function learning. Our results suggest that trial-by-trial analysis of behavioural data may provide unique insights into the BN phenotype, which may thus be computationally distinct from the related disorder of BED and the HC state.

## Introduction

Bulimia nervosa (BN) and binge eating disorder (BED) are eating disorders characterized by compulsive intake of food in excess quantities with a sense of lack of control over the episodes (***American Psychiatric Association, 2013***). In addition, patients with BN also exhibit inappropriate compensatory behaviours such as vomiting, laxative use, or excessive physical activity, to name a few. BED is the most common eating disorder, with a 12 month prevalence of 1.2%, compared to 0.3% 12-month prevalence for BN (***Hudson et al., 2007***). These disorders are often associated with comorbid psychopathology, including obsessive compulsive disorder, attention-deficit hyperactivity disorder, and alcohol or illicit drug dependence (***Hudson et al., 2007***).

The presentation and comorbidity patterns observed in pathological binge eating suggest dysfunction in the decision-making systems of afflicted individuals. This claim is supported by a growing body of empirical work which has implicated various putative components of the decision making apparatus, including value learning for both states (***Frank et al., 2011***) and actions (***Reiter et al., 2016***), exploration-exploitation balance (***Morris et al., 2015***), and integration of instrumental control systems (***Gillan et al., 2016***; ***Voon et al., 2015***). More specifically, eating disorder symptoms in a large online sample have been associated with decision making and learning strategies that rely on simply repeating previously rewarded actions (known as a habitual or model-free control) (***Gillan et al., 2016***). This contrasts with strategies that employ an internal model of the action-outcome contingencies in the environment in order to plan ahead (a so called model-based or goal-directed strategy). The ability to plan ahead facilitates flexible adaptation to a changing environment, since things that were once rewarding may at some point cease to be so. Impairment in model-based (MB) planning—with corresponding greater reliance on model-free (MF) control—has been shown in a clinical sample of patients with BED (***Voon et al., 2015***). Those subjects also showed increased reliance on value-agnostic strategies, namely perseveration, whereby they simply repeated their previous choices regardless of those actions’ expected values.

Less is known about the computational structure of decision making in BN. While prior work has evinced deficits in encoding critical value learning signals (***Frank et al., 2011***) and learning response biases toward rewarding stimuli (***Grob et al., 2012***), to our knowledge there is no characterization of control system integration in BN. Patients with BN often report difficulty overcoming the “force of habit.” That is their binge eating and purging behaviour becomes increasingly compulsive in nature (***Pearson et al., 2015***; ***Treasure et al., 2018***) which could speak to excessive reliance on MF control, similar to subjects with BED. Moreover, it is unclear how deficits in MB control are associated (if at all) with eating disorder symptom severity in a clinical sample. Finally, whether measures of MB and MF balance can differentiate subjects with pathological binge eating from each other and from controls is unknown.

The present study investigates the relationship between pathological binge eating diagnoses (BED & BN) and symptom severity—as measured by a commonly used eating disorders questionnaire‒ and the balance of MB and MF control in a clinical sample. We show that eating disorder symptom severity is associated with reduced MB control, but that BN (in contrast to BED) may be characterized by deficits in MF control. To this end, we demonstrate that BN can be separated from BED and HC primarily on the basis of MF system impairment, suggesting impaired value function learning in this group.

## Results

Ethical approval for this study was obtained from the Nova Scotia Health Authority (NSHA) Research Ethics Board and informed consent was obtained from all participants. Through online ads and the local adult eating disorders clinic, we recruited 25 healthy controls (HCs), 25 patients with BED, and 25 patients with BN for participation in the present study. ***Figure 1*** demonstrates our subject recruitment pipeline. An online pre-screening questionnaire that included the Eating Disorders Diagnostic Survey (EDDS) (***Stice et al., 2000***; ***Krabbenborg et al., 2012***; ***Stice et al., 2004***) was completed by 344 individuals. Of these individuals, 255 were interviewed by one of the investigators (ARK/AN) to ascertain satisfaction of inclusion and exclusion criteria (***Figure 1***). Included subjects were asked to visit our clinic for a single session consisting of (1) performance of the two-step task (***Daw et al., 2011***) (***Figure 2***), (2) questionnaire completion, and (3) completion of the operation span working memory task (OPSPAN; ***Conway et al. (2005)***; ***Unsworth et al. (2005)***; ***Otto et al. (2013)***).

Subject demographics and covariate summary statistics are presented in ***Table 1***. Distributions of subject demographics and rating scale data are displayed graphically in ***Appendix 1***. There were significant differences between groups across several covariates. The BED group was older, and demonstrated greater BMI. Pathological binge eating groups demonstrated higher scores on the obsessive-compulsive inventory-revised (OCI; ***Foa et al. (2002)***), and the Barratt impulsivity scale (BIS-11; ***Patton et al. (1995)***). Scores on the eating disorder examination questionnaire (EDE-Q; ***Fairburn and Beglin (1994)***) were significantly greater in the pathological binge eating groups. However, subjects with BED and BN were relatively evenly distributed amongst the higher scores on this questionnaire. Behavioural inhibition scores (BIS; ***Carver and White (1994)***) were highest amongst healthy controls. There were no significant differences in IQ (measured with the North American Adult Reading Test; NAART; ***Uttl (2002)***), working memory, or behavioural activation scores.

**Figure 1.**
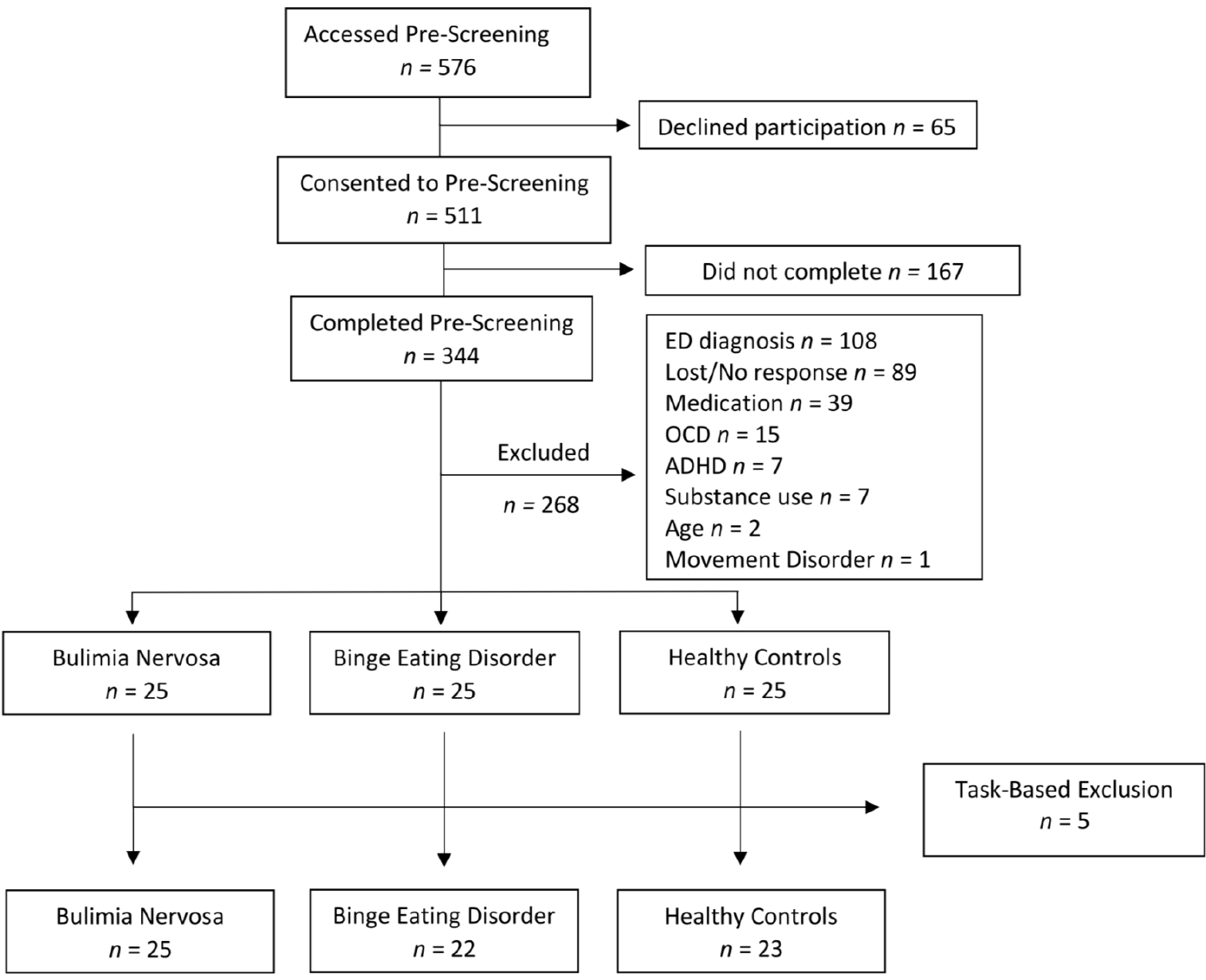
CONSORT flow diagram (www.consort-statement.com) demonstrating recruitment pipeline. We excluded 108 individuals who had disordered eating but did not meet DSM-5 criteria for bulimia nervosa or binge eating disorder. We excluded individuals who met DSM-5 criteria for obsessive-compulsive disorder (OCD; n=15), attention deficit-hyperactivity disorder (ADHD; n=7), substance use disorder (n=7), age over 40 years (n=2), and movement disorder (n=1). We excluded 39 subjects who were using either psychostimulants, dopamine D2-receptor antagonists, dopaminergic antidepressants such as bupropion or sertraline, other dopaminergic drugs such as L-DOPA or dopamine agonists, and serotonin-norepinephrine reuptake inhibitors (although venlafaxine at doses <150mg was not excluded). After clinical interview, we included 25 subjects with bulimia nervosa, 25 subjects with binge eating disorder, and 25 healthy controls. Healthy controls were free of any historical diagnosis of eating disorder. Five subjects were excluded based on task-related criteria: pressing the same key on more than 95% of trials (n=1), and making the same choice at the first step of the task on more than 95% of trials (n=4).

**Table 1.** Demographics and group characteristics. Statistics are either presented as means and standard deviation for normally distributed covariates (in round parentheses), or as median and interquartile range (in square parentheses). Abbreviations: healthy control (HC), binge eating disorder (BED), bulimia nervosa (BN), body mass index (BMI), obsessive compulsive inventory-revised (OCI; ***Foa et al. (2002)***), eating disorder examination questionnaire (EDE-Q; ***Fairburn and Beglin (1994)***), behavioural inhibition scale (BIS; ***Carver and White (1994)***), behavioural activation scale (BAS; ***Carver and White (1994)***), Barratt impulsivity scale (BIS-11; ***Patton et al. (1995)***), intelligence quotient (IQ; approximated with the North American adult reading test; ***Uttl (2002)***), and operation span (OPSPAN) score (***Conway et al., 2005***; ***Unsworth et al., 2005***).

|  | HC (n=23) | BED (n=22) | BN (n=25) | p |
| --- | --- | --- | --- | --- |
| Age | 24.00 [22.50, 30.00] | 33.00 [25.00, 36.75] | 22.00 [20.00, 30.00] | 0.001 |
| $n_{female}$ (%) | 23 (100.0) | 21 (95.5) | 24 (96.0) | 0.6 |
| BMI | 21.80 [21.19, 24.20] | 33.65 [29.33, 39.90] | 23.83 [21.70, 28.04] | <0.001 |
| OCI | 7.00 [3.50, 8.00] | 16.00 [10.25, 19.50] | 18.00 [13.00, 22.00] | <0.001 |
| EDE-Q | 0.39 [0.18, 0.54] | 3.66 [3.03, 4.75] | 4.22 [3.70, 4.75] | <0.001 |
| BIS | 13.00 [12.00, 16.00] | 12.00 [9.00, 14.75] | 10.00 [8.00, 12.00] | 0.001 |
| BAS | 25.87 (3.96) | 24.68 (6.07) | 24.68 (4.53) | 0.637 |
| BIS-11 | 55.09 (6.90) | 63.82 (8.38) | 65.92 (11.95) | <0.001 |
| IQ | 105.68 (9.33) | 105.33 (10.16) | 99.91 (8.84) | 0.066 |
| OPSPAN | 48.00 [34.50, 50.00] | 47.00 [27.25, 54.75] | 36.00 [24.00, 49.00] | 0.199 |

## Decision-Making Task and Theory-Free Analysis

The decision-making task employed was originally presented by ***Daw et al. (2011)*** (***Figure 2***). Subjects are initially (i.e. at the first-step) presented with two choices, from which they must select one. Each choice leads to one of two second-step states, with fixed probability. Once presented with the second-step choices, subjects must again choose one of the two options, after which a reward is either presented or omitted. The probability of reward for each of the 4 second-step choices is independent of the others, and varies on a trial-by-trial basis (the probabilities always lie between 0.25 and 0.75; ***Daw et al. (2011)***). Included subjects performed 201 trials in 3 blocks of 67 trials, within the same testing session.

To probe the effects of past trial behaviours and their outcomes on subsequent decisions, we implemented a convolutional logistic regression previously described by ***Miller et al. (2016)*** and similar to that implemented by ***Lau and Glimcher (2005)***, which we simplify graphically in ***Figure 3***. This analysis is based on the fact that the two-step task contains eight possible trial events. These include trials with a reward after common transition (RC), reward after an uncommon transition (RU), reward omission after a common transition (OC), and reward omission after an uncommon transition (OU); for each of these, the subject may have chosen action A or B at the first step. A subject’s behavioural data for a set of ***T*** trials can be represented as a ***T*** × 1 vector of choices y ∈ {0,1}^*T*×1^ (where for example 1 = Choice A, and 0 = Choice B at the first step), and a **T** × 4 matrix 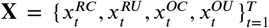 identifying the type of trial encountered at timestep *t*. Using logistic regression, we can learn a filter of weights 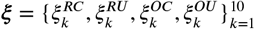 over a 10-trial window, where, for example 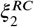 represents the influence of having experienced trial *x*^*RC*^, 2 trials ago. We can represent this logistic regression compactly as

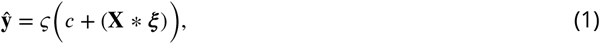

where **ŷ** is a vector of predicted action probabilities for trials ***t* =** {11,… **,*T***}, **ζ** is the sigmoid function, **c** is an intercept term, and **X** * ξis the convolution of **X** and ξ (without zero padding). Assuming the parameters (c, ξ) for each subject vary around a group-level mean and standard deviation, we perform inference using hierarchical linear mixed effects modeling (using `lme`4 package in the R statistical programming language; ***Bates et al. (2015)***). ***Figure 3b*** and ***Figure 3c*** demonstrate filters learned for hypothetical MB and MF learners, respectively, illustrating that the shape of filter ξ can capture a subject’s control strategy.

**Figure 2.**
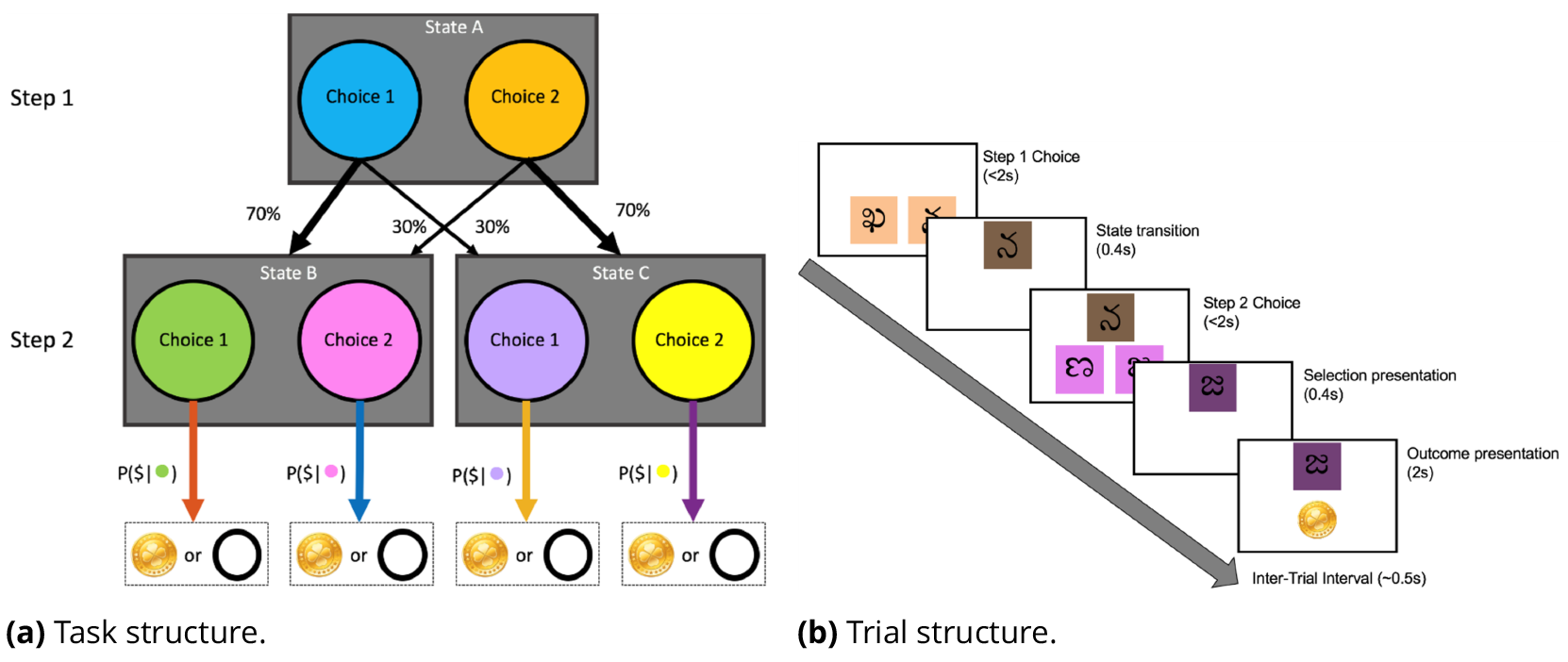
Individuals are initially presented with the first step that contains two options, from which subjects must select one by pressing a key. Choices must be made within 2 seconds of stimulus presentation. Selection of a first-step option will lead to a given second-step state with probability of 70% (the “common” transition), and to the other with a probability of 30% (the “rare” or “uncommon” transition;duration of transition is 0.4s). These probabilities remain fixed throughout the experiment. Each of the two second-step states has nested within it a set of two options from which the subject again selects by pressing a key (time limit is 2 seconds); those second-step state options are each associated with a reward probability that varies between 0.25 and 0.75 according to a Gaussian random walk with standard deviation 0.025. The image selected at Step 2 then moves to the top of the screen for 0.4 seconds at which time feedback is provided in the form of a reward (gold coin image) or no reward (empty circle).

By comparing the parameter estimates *ξ* for each trial back, we can evaluate the degree of MB and MF control reflected in a subject’s data; we denote these as the model-based index (MBI) and model-free index (MFI). We found a negative association of symptom severity—as measured by EDE-Q—with MBI (Pearson’s r=−0.43;p<0.001) but not MFI (***Figure 4a***). The association between EDE-Q and MBI remained statistically significant after correction for BMI and OCI-R, BIS, BAS, BIS-11, IQ, and OPSPAN scores (regression coefficient −4.48, p=0.011;***Table 2***), while the relationship between EDE-Q and MFI remained not statistically significant (2.12, p=0.085). Further, we submitted MBI and MFI as features to a sparse linear classifier whose target output was diagnostic group (in a one-vs-rest format). Subjects with BN were accurately discriminated from BED and HC with an area under the receiver operating characteristic curve (AUC) of 0.78 (95% CI = 0.64-0.92; ***Figure 4b***). Inspection of the MBI and MFI values (decomposed over each “step back” in the filter *ξ*) shows a trend toward lower values of both MBI and MFI in subjects with BN (***Figure 4c***), although the sparse linear classifier weights highlight MFI at the previous trial as the most important factor underlying classification performance (***Figure 4d***).

## Analysis of Reaction Times

Like previous studies (***Deserno et al., 2015***),we collected reaction time data from participants during performance of the two-step task. We were particularly interested in the second-step reaction time as a proxy measure for the degree to which the subject learned the state-transition model. Greater reliance on the MB system involves use of estimated transition probabilities in first-step choices; in subjects using such a strategy, rare transitions are assumed to elicit surprise, which this analysis component assumes will slow reaction time at the second step. Indeed, for every group, the MBI was positively associated with reaction time slowing after rare transitions (***Figure 5***). This correlation was strongest for HC subjects (r=0.82, p<0.001), followed by both BN and BED groups (r=0.77, p<0.001 for each). Only the BN group demonstrated a statistically significant correlation between the MFI and reaction time difference (r=0.57, p=0.0029).

**Figure 3.**
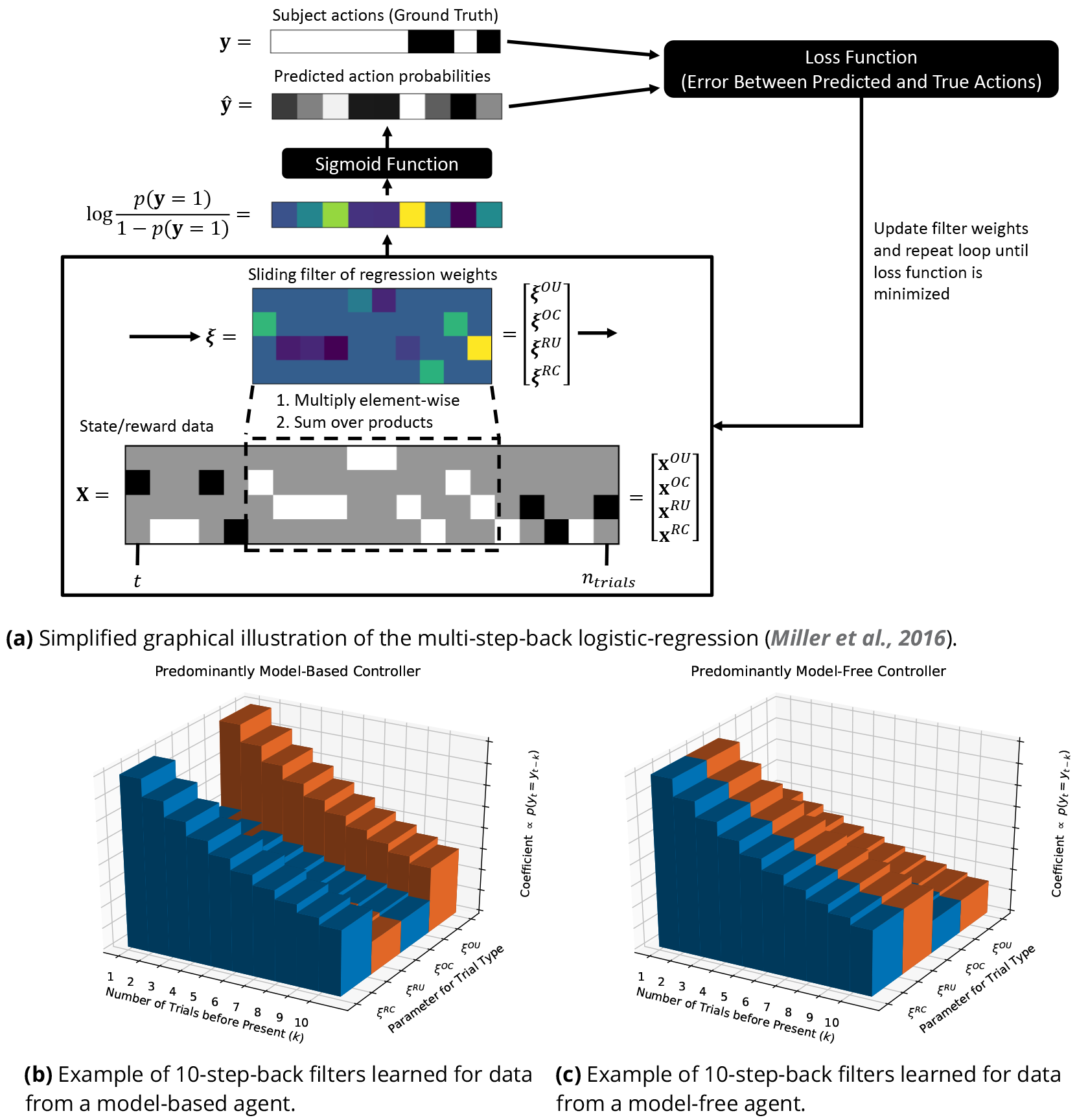
***Figure 3a*** shows the iterative process of learning filter weights for a single subject. The analysis done in our study modeled filter weights as random variables distributed around a group-level mean. ***Figure 3b*** and ***Figure 3c*** illustrate the filter values that would be learned for model-based and model-free agents, respectively. The filter value is proportional to the probability that a subject will repeat an action taken at a given number of trials before the present, for a given trial type.

**Figure 4.**
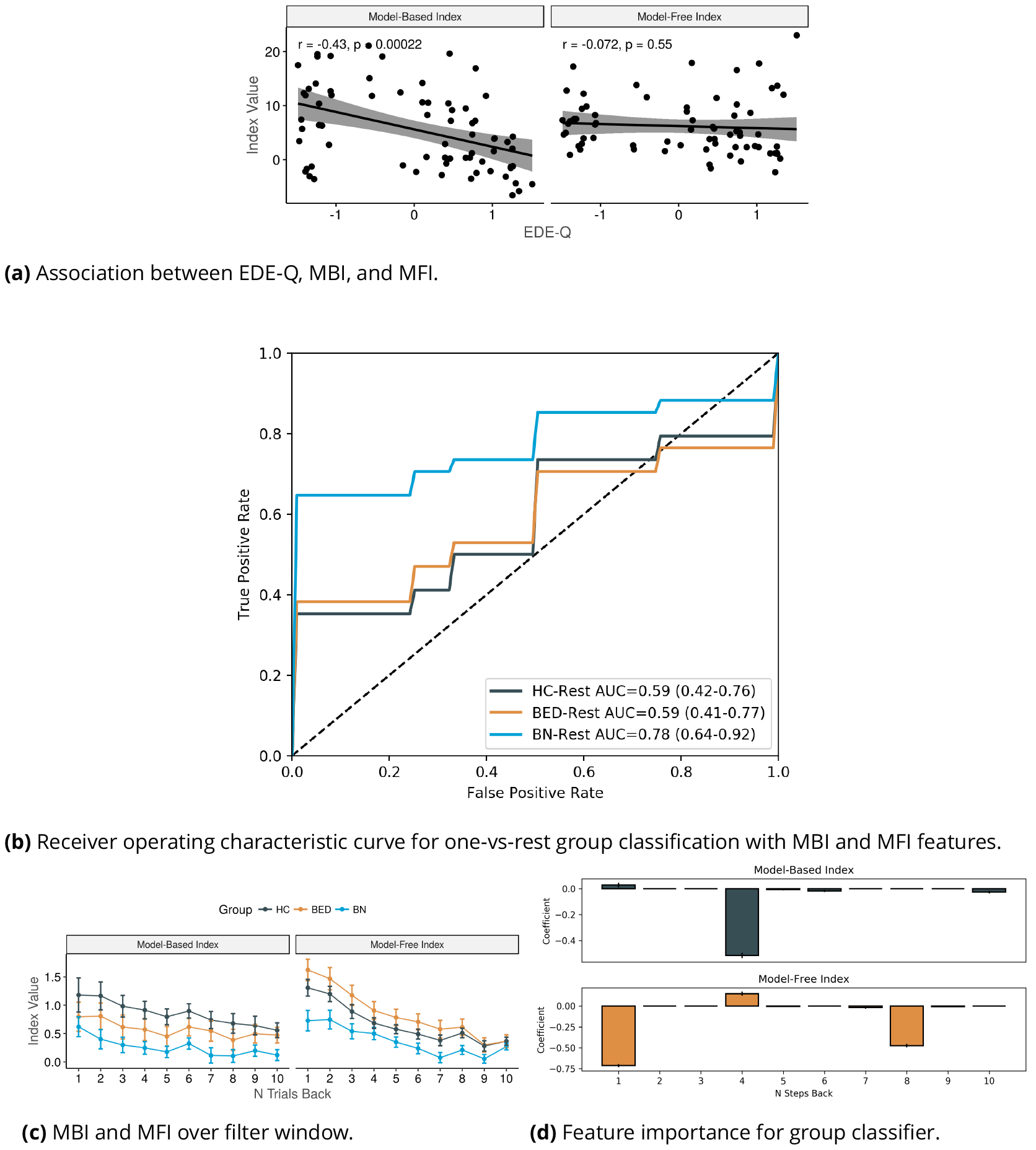
***Figure 4a:*** Relationship between eating disorders examination questionnaire (EDE-Q) score and the MBI, MFI estimates. There was a statistically significant negative correlation between EDE-Q and MBI (r=−0.43, p<0.001). This relationship remained significant after correction for covariates (p<0.001;see ***Table 2***). Similar results were shown for EDE-Q against estimates of model-based and model-free control parameters from reinforcement learning modeling. ***Figure 4b:*** Receiver operating characteristic (ROC) curve for sparse one-vs-rest logistic regression classifier of group based on MBI and MFI across 10 trials. Curves are coloured by the group being classified, and represent averages of ROC curves computed over each of 17 stratified cross-validation folds. BN subjects were the best classified (AUC 0.78;95% CI = 0.64–0.92). ***Figure 4c***: Mean MBI (left facet) and MFI (right facet) by group. The x-axis represents indices for the number of trials prior to the reference choice (i.e. “N Trials Back“). MBI and MFI values are plotted on the y-axis. Error-bars are standard errors. Lines are coloured by group: healthy controls (HC), binge eating disorder (BED), and bulimia nervosa (BN). ***Figure 4d***: Sparse logistic regression coefficients averaged across 17 folds of stratified cross validation. The upper plot shows coefficients for the model-based index (MBI) over each step prior to a given trial. The bottom row shows coefficients for the model-free index over the same period. The x-axis represents the number of trials before a given choice. The y-axis is the value of the regression coefficient. Error bars are standard errors. MFI at the previous trial was the most important feature.

**Table 2.**
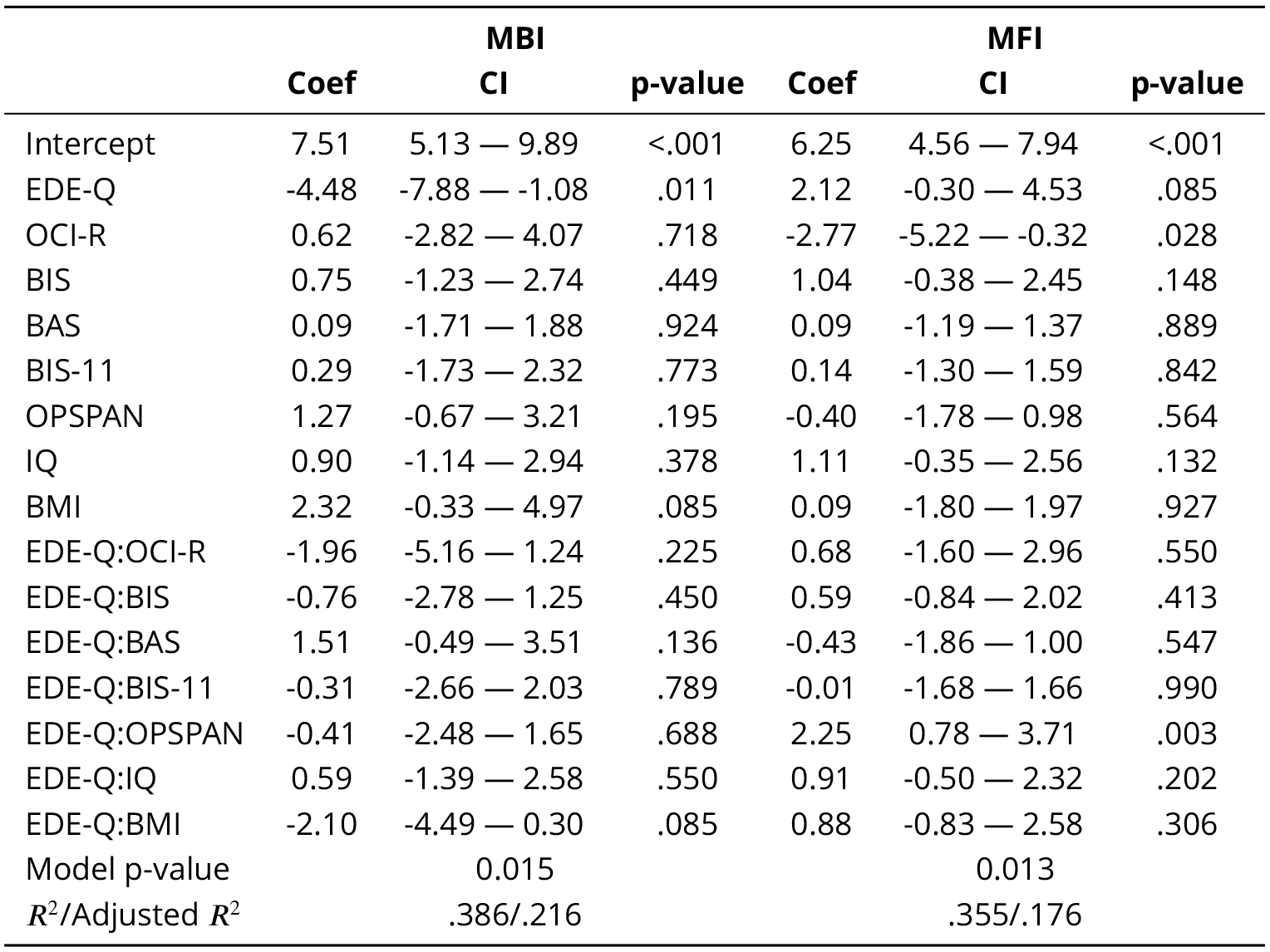
Coefficients of linear regression between MBI, MFI, and questionnaires. Abbreviations: 95% confidence interval (CI), model-based index (MBI), model-free index (MFI), eating disorder examination questionnaire (EDE-Q), obsessive-compulsive inventory revised (OCI-R), behavioural inhibition scale (BIS), Barratt Impulsivity Scale (BIS-11), operation span (OPSPAN).

**Figure 5.**
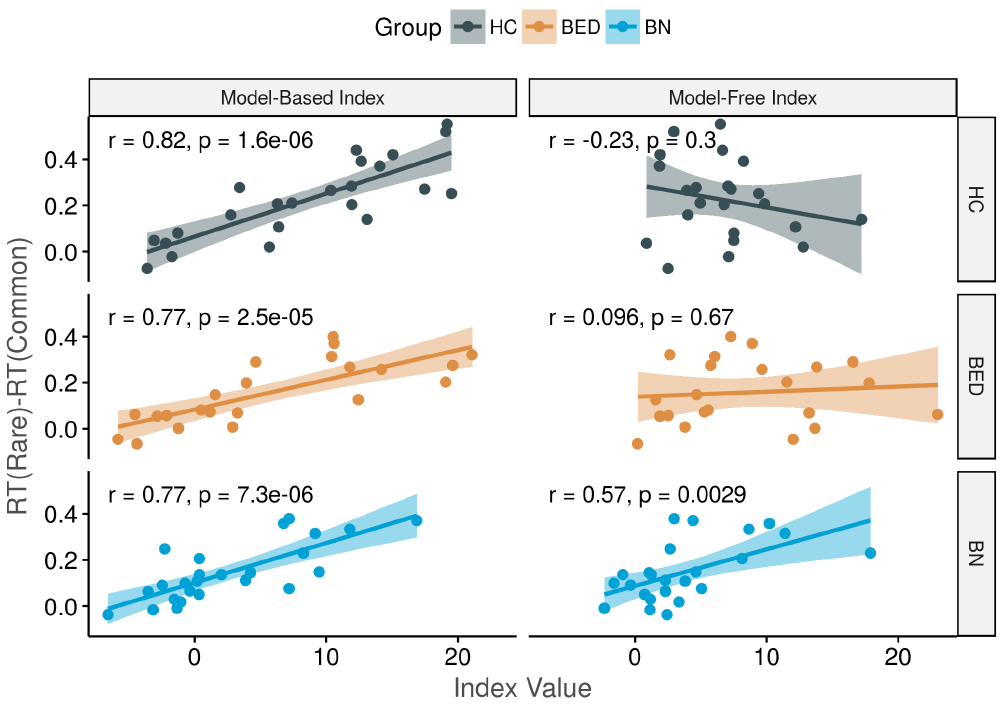
Second step reaction time differences against MBI and MFI, faceted by group. The x-axis shows the value of model-based index (left column) and model-free index (right column), and the y-axis plots the difference between reaction time after rare and common transitions, respectively. Each row plots this relationship for a given group: healthy controls (HC), binge eating disorder (BED), and bulimia nervosa (BN). The model-based index is positively associated with reaction time difference for each diagnosis, and these associations were all statistically significant. The model-free index was associated with reaction time difference only in the BN group.

**Figure 6.**
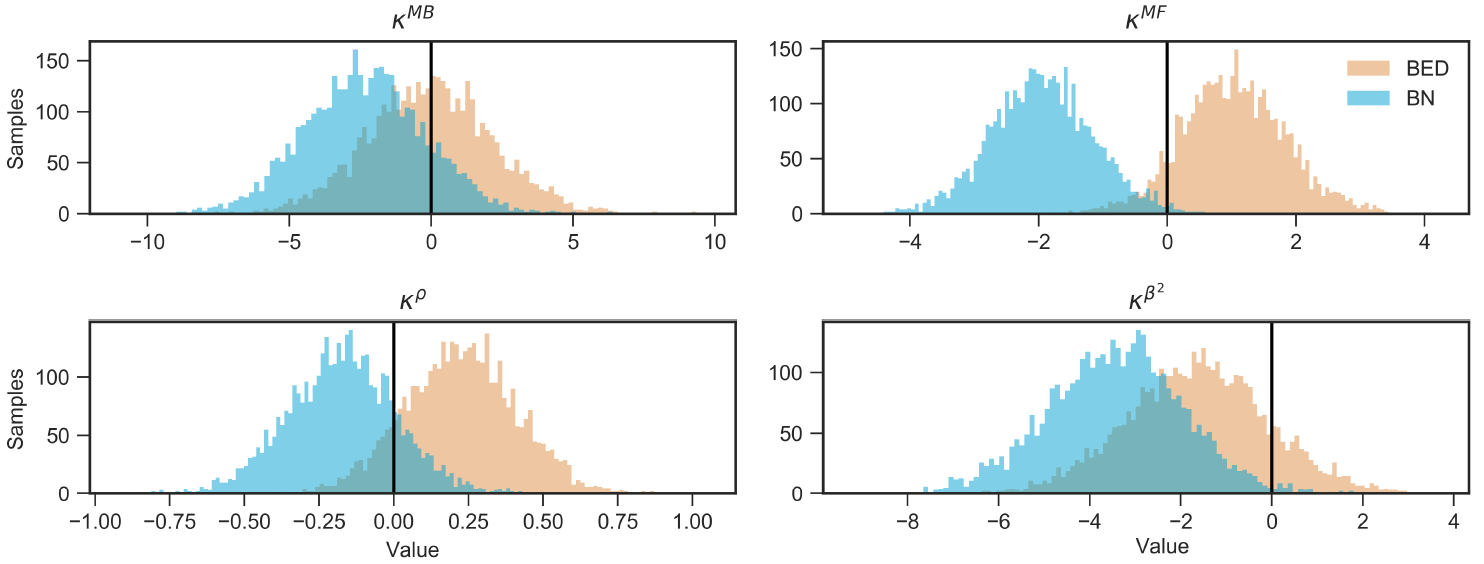
Generalized linear model coefficients from the reinforcement learning model fit. Plots are histograms of samples from the posterior probability density over the **κ** parameters. The x-axis shows the **κ** parameter value, and the y-axis represents the count of samples over that value. Histograms are coloured by group. Note these parameters represent the effect of group (BED or BN) on the respective parameter estimate (e.g. ***β*^*ω*^** for ***κ***^***M B***^ in this Figure) above the level expected from healthy controls. In general, bulimia nervosa showed lower values of all inverse softmax temperatures relative to healthy controls. These differences were either greater than those observed in the binge eating disorder group, or opposite in direction.

## Reinforcement Learning Modeling

To further investigate the findings of theory-free analyses, we fit an RL model to subjects’ behavioural data. We implemented a computational model similar to that in ***Sharp et al. (2015)***. Each subject was characterized by a set of parameters 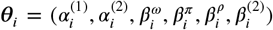, where 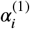,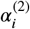 are learning rates at the first and second step, respectively, 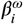 is a parameter describing the influence of MB state-action values on action selection, and 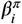,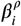 are similar weights for MF state-action values and perseveration, respectively. We included a separate weight for values at the second-step, 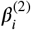, since our separation of MB and MF control is focused mainly on Step 1 choices. Group effects (HC, BED, BN) on 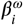, 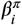, 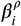, 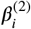 were estimated jointly during model fitting in the same fashion described by ***Sharp et al. (2015)***. For example, the effect of BED diagnosis (with respect to HC diagnosis) on the **ω** parameter would be denoted as 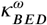, and the interpretation is similar to that of ordinary generalized linear model weights.

We show estimates of the generalized linear model coefficients in ***Table 3***, and plot posterior samples in ***Figure 6***. We found that a reduction in MF control was the primary difference between subjects with BN and healthy controls (mean 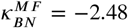 95% CrI −3.53 to −0.38; where CrI is a credible interval). Patients with BN also showed lower values of ***β***^(2)^ (mean 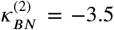 95% CrI −6.46 to −0.62), suggesting noisier decision making at the second step of the task. The credible intervals for group effects on model-based control, ***κ^*M B*^*** included 0 for both BN (95% CrI −6.62 to 1.61) and BED (95% CrI −4.49 to 4.03), as did the model-free effects for BED (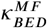 95% CrI −0.58 to 2.79). The correlation between ***β*^*ω*^** (MB control parameter) and EDE-Q was statistically significant (correlation coefficient r=−0.25, p=0.038), although correlation of MF control with EDE-Q was not (r=—22120.17, p=0.15); this is in line with the main effect shown with the theory-free analysis.

Projection subjects onto the first three principal components of MBI, MFI, RL model parameters, and the reaction time difference is shown in ***Figure 7***. We expected that theory-free MBI and MFI would align with 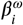 and 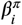, respectively, and that the reaction time slowing would align predominantly with 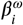, which were supported by the data. MFI was roughly orthogonal to the model-based control measures on the first 2 principal components. Notably, there was no overlap in the 95% confidence interval (represented by the ellipse) of BN’s projection with those of the other groups. Moreover, the BN subjects are projected onto a region associated with lower values of nearly all parameters.

**Table 3.** Summary of posterior distribution over group-level effects on reinforcement learning (RL) parameter estimates. Each ***κ*** is the coefficient of a generalized linear model defining the effect of being a HC, BN, or BED subject on the consequent RL parameter estimate. For a given parameter *x*_**i**_ for subject *i*, its distribution is 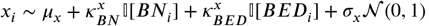 where 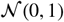 is a standard normal variate, and *μ*_***x***_, *σ*_***x***_ are the mean and standard deviation for parameter ***x*** over all subjects. We show the mean and percentiles for each of these estimates. The 95% posterior credible interval is the interval between the 2.5 and 97.5 percentiles of posterior samples.

| Parameter | Mean | Percentiles |  |  |  |  |
| --- | --- | --- | --- | --- | --- | --- |
|  |  | 2.50% | 25% | 50% | 75% | 97.50% |
| $\kappa_{BN}^{MB}$ | -2.48 | -6.62 | -3.89 | -2.52 | -1.04 | 1.61 |
| $\kappa_{BN}^{MF}$ | -1.97 | -3.53 | -2.53 | -1.99 | -1.42 | -0.38 |
| $\kappa_{BN}^{\rho}$ | -0.17 | -0.54 | -0.3 | -0.17 | -0.04 | 0.18 |
| $\kappa_{BN}^{\beta^{(2)}}$ | -3.5 | -6.46 | -4.48 | -3.47 | -2.53 | -0.62 |
| $\kappa_{BED}^{MB}$ | -0.2 | -4.49 | -1.67 | -0.18 | 1.28 | 4.03 |
| $\kappa_{BED}^{MF}$ | 1.06 | -0.58 | 0.48 | 1.05 | 1.63 | 2.79 |
| $\kappa_{BED}^{\rho}$ | 0.24 | -0.17 | 0.11 | 0.24 | 0.37 | 0.63 |
| $\kappa_{BED}^{\beta^{(2)}}$ | -1.66 | -4.65 | -2.65 | -1.67 | -0.7 | 1.46 |

**Figure 7.**
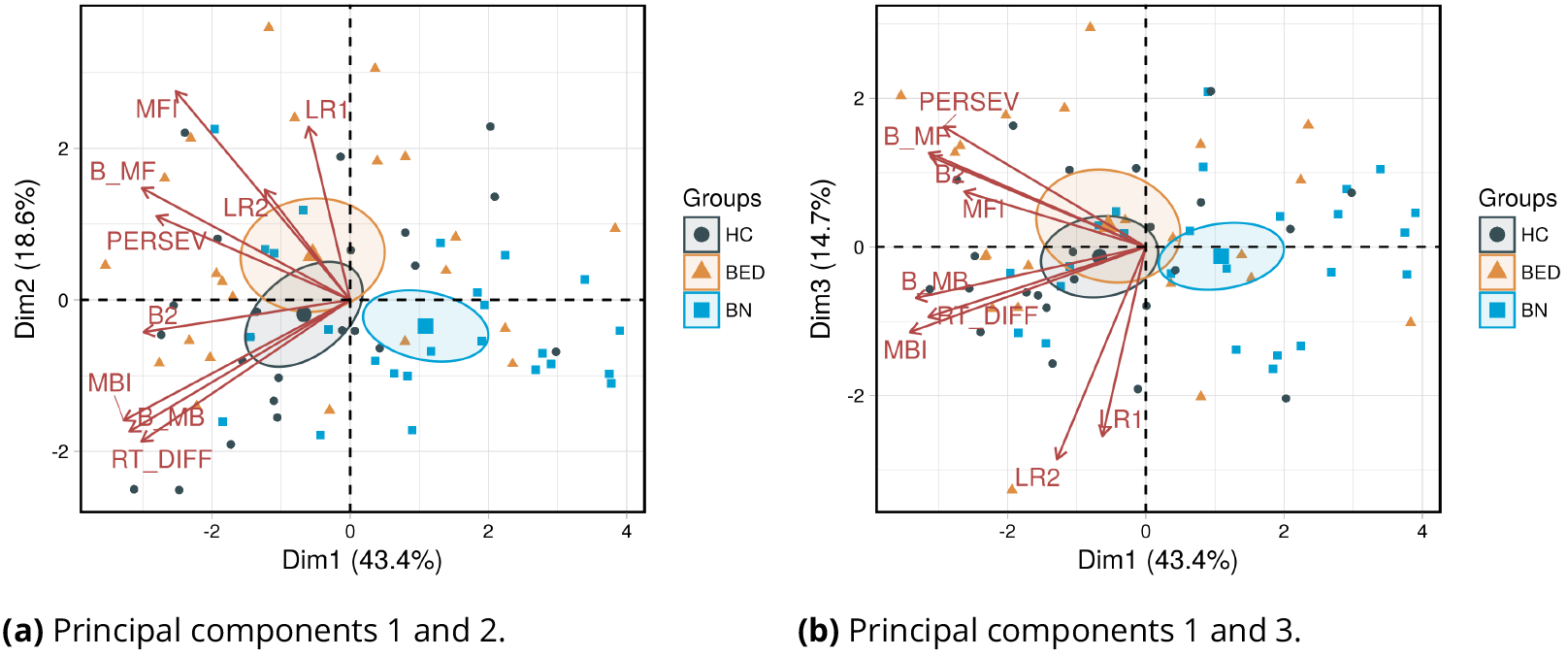
Principal components analysis of parameters estimated in theory-free analysis and RL modeling. ***Figure 7a*** shows data projected onto the first two principal components, while ***Figure 7b*** shows data projected onto principal components 1 and 3. Ellipses are 95% confidence intervals about the mean (markers centered in ellipsoids) of data for each group. Abbreviations (“; RL” in parentheses indicates parameter was from RL model): model-based weight (B_MB;RL), model-free weight (B_MF;RL), perseveration parameter (PERSEV;RL), step-2 inverse softmax temperature (B_2;RL), step-1 learning rate (LR1;RL), step-2 learning rate (LR2;RL), reaction time difference between rare and common trials (RT_DIFF), model-based index (MBI), model-free index (MFI).

## Discussion

We have shown that increased eating disorder severity, as indexed by the EDE-Q, is associated with impaired MB control in healthy controls and a clinical sample of patients with BN and BED, although severity was independent of MF control. Our results also suggest that impairments in MF control may be a prime deficit in patients with BN. Importantly, we showed that the MF deficit was of sufficient magnitude to discriminate BN subjects from other groups in our study. In this section, we posit that failure to learn the environmental value function may be a characteristic impairment of the BN reinforcement learning phenotype.

Our first finding was of an inverse relationship between MB control and eating disorder symptom severity. This result contributes to those of ***Gillan et al. (2016)***, who previously showed impairments in MB control associated with higher scores on a scale commonly used for eating disorder screening (***Grob et al., 2015***). Those authors found that questionnaires related to alcohol use and obsessive-compulsive symptoms were also associated (through a common factor) with reduced MB control. Our paper addressed these potentially confounding factors by carefully screening out subjects with substance use. It was difficult to screen out obsessive-compulsive symptoms in the clinical participants to the levels present in the healthy controls, and so we included a measure of these symptoms as a covariate in our analysis. That the relationship between EDE-Q score and lower MB control withstood control for covariates suggests that eating disorder symptomatology may independently affect the degree of MB control used. Our clinical sample included currently ill individuals, so it is possible that the nutritional changes in the midst of BED or BN may account for this excess MB control impairment. Future investigations could address this through within-subjects designs across ill and remitted states. To our knowledge, it is unclear whether impaired MB control is a state or trait phenomenon in eating disorders.

Interestingly, our data suggest that BED and BN are associated with different balances of MB and MF control. For subjects with BED, there was a tendency toward greater use of MF control compared to HCs, although the 95% credible interval did not exclude 0 (Figure 5 and Table 2; 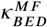 95% CI −0.58 to 2.79). This finding, as expected, was in line with that of ***Voon et al. (2015)***, who found greater reliance on habitual control in a clinical sample of patients with BED. It is possible that our study failed to conclusively replicate those results due to small sample size. Notwithstanding, the strongest statistical effect was observed for reduced MF control in BN subjects. This was supported by accurate discrimination of BN from both BED and HCs using a sparse linear classifier (ROC-AUC 0.78, 95% CI 0.64 −0.92) trained on indices of MB and MF control. Indeed, MF impairments were most important in this respect. This result was further substantiated by the results of RL modeling, which showed substantially impaired MF control in BN (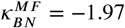; 95% CI-3.53 to-0.38). Failure to learn the environmental value function can explain these findings, and our data raise the possibility of two underlying computational mechanisms.

First, BN subjects may have reward insensitivity. Inability to sense reward would attenuate the target component of the reward prediction error signal, which drives value function learning. Unfortunately, the experimental paradigm employed herein is not suitable for estimation of reward sensitivity, so no direct evidence to this end is available. However, the literature supports the idea that pathological binge eating may result from reward hyper-and hypo-sensitivity in BED and BN, respectively (***Friedrich et al., 2013***). Under a Pavlovian conditioning paradigm, ***Frank et al. (2011)*** demonstrated impaired temporal difference reward prediction error signaling in subjects with BN, compared to healthy controls—a deficit related to the frequency of binging and purging symptoms. Dopaminergic tone may underlie these effects, as demonstrated by ***Grob et al. (2012)***, who exposed 19 subjects with remitted BN to catecholamine depletion and found a greater impairment in reward responsiveness compared to healthy control subjects. Experimental catecholamine depletion has also been shown to increase the expression of bulimic symptoms in patients with remitted BN (***Grob et al., 2015***). Thus, a primary impairment in sensing the reward required to learn a task value function could bring about our observations in BN patients, and this could be related to both dopaminergic signaling and clinical symptoms.

The results of both theory-based and theory-free models may have been explained by excessively noisy action selection, which could arise either (A) through excessive but directed exploration, or (B) by sheer random choice. Under the RL models posited herein, learning a value function requires relative stability in a behavioural policy, particularly at the second step. This is because the target element of the temporal difference prediction error, i.e. 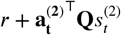, depends on the stability of the state action value computed at Step 2. To this end, our data showed greater step-2 choice randomness in BN (mean 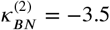 95% CI −6.46 to −0.62; also see the distribution of subjects in the PCA), which would not relate to subjects’ ability to learn the task transition function; reaction time data further suggested that BN subjects were able to learn state transition probabilities, since slowing was observed after surprising outcomes. To this end, a high choice randomness at the second step results in an inconsistent behavioural policy and learning becomes difficult because of target instability. Indeed this has been a well-known problem in the artificial intelligence literature, wherein important measures to maintain target stability are common (***Sutton and Barto, 1998***; ***Mnih et al., 2015***). Future work should focus on disentangling the roles of choice stochasticity and reward sensitivity (or reward learning rate) of the RL systems in subjects with BN.

A strength of the present study is the use of a more robust theory-free analysis method (***Miller et al., 2016***) to identify effects for subsequent reinforcement learning modeling. Another strength of our study is the presentation of classification performance using subject-level parameters inferred from models. By this application, we emphasize the focus on predictive validity in computational psychiatry. However, this predictive analysis was limited by our relatively small sample size. It would be of great interest to replicate the present analysis with a larger sample to determine whether the predictive capacity of MBI and MFI show resilience, and whether BED can be separated from HC and BN.

Our small sample size may have precluded a well-powered multi-label classification analysis, for which we substituted a one-vs-rest approach. Another limitation is that diagnoses were not made using gold standard research criteria such as the Structured Clinical Interview for DSM-5 (SCID-5) (***First et al., 2015***). However, the diagnostic tool employed in this study, the Eating Disorder Diagnostic Scale, was developed using DSM criteria and has adequate psychometric properties (***Stice et al., 2000***; ***Krabbenborg et al., 2012***; ***Stice et al., 2004***). The levels of agreement between EDDS and structured clinical interviews (including the SCID), as measured by the kappa coefficient, have been reported as .78-.91 for BN diagnoses and .72-.74for BED diagnoses, suggesting a high degree of agreement between these tools (***Stice et al., 2000***; ***Krabbenborg et al., 2012***; ***Stice et al., 2004***). Additionally, eating disorder diagnoses were confirmed by a psychiatrist with more than 10 years of experience working with these populations. Moreover, it is also possible that subjects recruited for the BED and BN populations may be more representative of a natural clinical sample, since we observed EDE-Q global scores consistent with reported norms from treatment-seeking patients (***Aardoom et al., 2012***; ***Welch et al., 2011***). However, no studies to our knowledge have compared differences between naturalistic samples of eating disorder patients with those classified by the SCID or other structure interviews.

In sum, we showed that greater eating disorder symptom severity is associated selectively with a reduction in MB (goal-directed) control, and that this effect may be independent of covariates known to be related with impaired MB control. However, in terms of ability to classify subjects based on diagnosis, impairments in MF control proved characteristic of subjects with BN. Our study is the first such investigation in a clinical sample of patients with both BN and BED, and highlights an important computational difference between these diagnoses.

## Methods and Materials

### Subjects

We recruited 75 participants (25 healthy controls, 25 BED, and 25 BN subjects). A sample size of 75 was computed as offering a power of 0.8 to detect correlation magnitude (of EDE-Q against model-based index) as low as 0.32 at a significance threshold of **α** = 0.05, while respecting our budgetary constraints. Power calculations were done in the pwr package for the R statistical programming language (https://github.com/heliosdrm/pwr). Clinical participants met DSM-5 (***American Psychiatric Association, 2013***) criteria for BED or BN, but were excluded if duration of illness was longer than 10 years. Prospective healthy controls were excluded if historically afflicted with any DSM-5 eating disorder(s). Clinical exclusion criteria for all subjects included history of obsessive-compulsive disorder, substance use disorder, depressive disorder, attention-deficit hyperactivity disorder, bipolar disorder, primary psychotic disorders, or movement disorders. Subjects were excluded if currently using psychostimulants, dopamine D2-receptor antagonists, dopaminergic antidepressants (bupropion, sertraline), other dopaminergic drugs (such as L-DOPA), or serotonin-norepinephrine reuptake inhibitors at doses greater than 150mg. Subjects were also excluded if a first-degree relative was afflicted with psychotic illness. Subjects were provided with $20 (CAD) in compensation for travel and parking, in addition to a token amount dependent on task performance. We excluded five subjects due to (A) four having selected the same first step option on more than 95% of trials, and one having pressed the same key on more than 95% of trials.

### Questionnaires

Subjects completed the following rating scales digitally: the behavioural inhibition scale/behavioural activation scale (BIS/BAS) (***Carver and White, 1994***), the Obsessive-Compulsive Inventory-Revised (OCI-R) (***Foa et al., 2002***), the Barratt impulsivity scale (BIS-11) (***Patton et al., 1995***), and the Eating Disorder Examination Questionnaire (EDE-Q) (***Fairburn and Beglin, 1994***). Intelligence quotient (IQ) was approximated using the North American Adult Reading Test (***Uttl, 2002***) administered by the investigators.

Continuous demographic and rating scale data were tested for normality using the Shapiro-Wilk test. Normally distributed data were summarized with mean and standard deviation, and differences between the three groups tested for statistical significance using analysis of variance. Non-normally distributed continuous variables were summarized as median and interquartile range (25%–75%), and tested for significance using the Kruskall-Wallis test. Categorical variables were summarized as absolute counts and proportions, and compared using Fisher’s exact test. A significance threshold of **α** = 0.05 was chosen a priori for all frequentist tests. Unless otherwise specified, all frequentist tests were two-sided.

### Decision-Making Task

We translated a two-step sequential decision-making task originally presented by (***Daw et al., 2011***) into the PsychoPy framework (v.1.84.0rc4; (***Peirce, 2007***)). In this task, subjects are initially (i.e. at the first-step) presented with two choices, from which they must select one. Each choice leads to one of two second-step states, with fixed probability. Once presented with the second-step choices, subjects must again choose one of the two options, after which a reward is either presented or omitted. The probability of reward for each of the 4 second-step choices is independent of the others, and varies on a trial-by-trial basis according to a Gaussian random walk with standard deviation 0.025, and bounded between 0.25 and 0.75 (***Daw et al., 2011***). Subjects were provided with training regarding the structure and dynamics of the task as in the original paradigm (***Daw et al., 2011***). Subjects performed 201 trials in 3 blocks of 67 trials, within the same testing session. Due to resource constraints, we shortened the trial step duration (primarily at transition points; Figure 1).

Task-based exclusion criteria included missing more than 10% of trials, demonstrating a mean reaction time >2 standard deviations faster than the overall group mean, responding with the same key on the keyboard for >95% of trials, and making the same choice for >95% of trials.

### Working Memory Task

Working memory was measured using the operation span task (OPSPAN) (***Unsworth et al., 2005***; ***Otto et al., 2013***). This task was implemented in PsychoPy (***Peirce, 2007***) according to a previous specification by ***von der Malsburg (2015)***. We implemented three iterations, each including spans between 3 and 7 consonant-equation pairs long. The upper limit of 7 was chosen based on internal piloting of the paradigm suggesting that 7 avoided ceiling effects. The OPSPAN score was calculated according to (***Unsworth et al., 2005***).

### Theory-Free Analysis of Behavioural Data

Here, we describe the details of the 10-step back theory free analysis (***Miller et al., 2016***; ***Lau and Glimcher, 2005***). Let 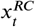 denote the occurrence of a common transition, C, and reward, *R* (hence “***RC***” superscript) at trial ***t***. If the *RC* combination of transition and reward occurred after choosing option A (Choice 1 in ***Figure 2a***) at the first step, then let 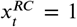; conversely, had the *RC* event occurred after selection of option B (Choice 2 in ***Figure 2a***), then let 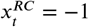. If the event *RC* did not occur at trial t, then let 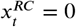. Thus, a subject’s behavioural data for a set of *T* trials as a *T* x 1 vector of choices y (where for example 1 = Choice A, and 0 = Choice B at the first step), and a *T* x 4 matrix

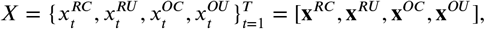

where superscripts are the events RC (Rewarded, Common transition), RU (Rewarded, Uncommon transition), OC (reward Omitted, Common transition), and OU (reward Omitted, Uncommon transition). Note that each row of **X** is zero everywhere except the column denoting the last trial’s event.

If at trial ***t*** − 1 one observes 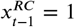 (which would equivalently be captured as *x*_***t***-1_ = [1,0,0,0], and subsequently the subject makes a first step choice of *y*_**t**_ = 1, we are interested in describing the influence of 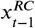. Introducing the parameter ξ^**RC**^ and denoting 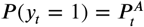 we can roughly define a linear relationship between the log odds of Choice A and 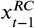:

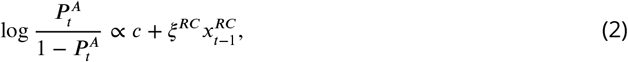

where *c* is a scalar intercept parameter capturing an independent tendency to select Choice A. letting *ξ*^***RC***^ be a *K* × 1 vector 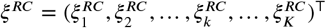 allows us to identify the influence of observation 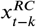 which occurred *k* steps before the present trial t. The relationship between trial type *RC* on the first step choice at trial *t* can thus be summarized as

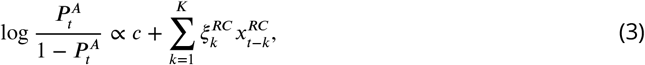

which is easily extended to other trial types (RU, OC, OU), whence the following regression equation is proposed ***Miller et al. (2016)***:

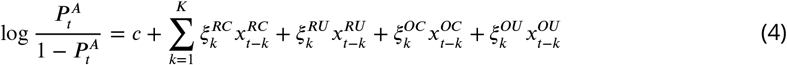

From Equation ***Equation 4***, we derive a model-based index (MBI) and model-free index (MFI), which provide measures of the respective use of MB and MF strategies ***Miller et al. (2016)***:

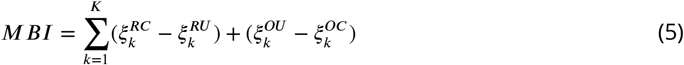

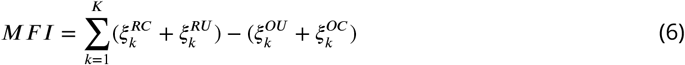

We performed inference of ***ξ*** and ***c*** across subjects using generalized linear mixed effects modeling with the lme4 package in R (***Bates et al., 2015***). We begin with a dataframe object with the following columns:

- y (int) ∈ {0,1}, representing action taken at each trial (indexed by `trial_number`)
- `subject_id` (Factor)
- trial_number (int)
- RC (int ∈ {−1,0,1}) indicating whether trial `trial_number-k_back` was of RC type
- RU (int ∈ −1,0,1) indicating whether trial `trial_number-k_back` was of RU type
- OC (int ∈ −1,0,1) indicating whether trial `trial_number-k_back` was of OC type
- OU (int ∈ −1,0,1) indicating whether trial `trial_number-k_back` was of OU type
- k_back(int) ∈ {1,2,…, 10}

To estimate a matrix ξ ∈ ℝ^κ×4^ of parameters and an intercept *c*_*i*_ for the *i*^*th*^ subject, we implemented the following regression model (in R notation for lme4 package):

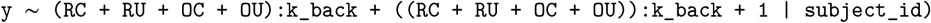

After estimating ***ξ*** and ***c*** for each subject, we compared MBI and MFI with EDE-Q using linear regression to determine the effect of symptom severity on control indices; this model was corrected for variation in other covariate scales including OCI-R, BIS, BIS-11, and the working memory score from the OPSPAN.

The global EDE-Q score was the primary variable of interest in terms of relationship with MB and MF control. This is a 28-item self-report measure of four domains including eating concern, weight concern, shape concern, and dietary restraint over the past 28 days. The measure is adapted from the widely used Eating Disorder Examination interview. The literature provides support for the reliability and validity of the EDE-Q for measuring eating disorder symptoms among adult women (***Fairburn and Beglin, 1994***). In the present study, the measure demonstrated excellent internal consistency (Cronbach’s **α** = .98).

### Group Classification with Sparse Linear Classifier

The purpose of this analysis was to determine whether the filters learned in the multi-step back logistic regression offered discriminative power with respect to diagnosis. For a given subject we define 1 × ***K*** vectors, **m**^*M F I*^ and **m**^*M B I*^, which are

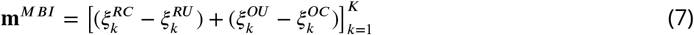

and

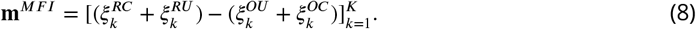

Given a sample size of *N*, we then construct an ***N*** × 2***K***

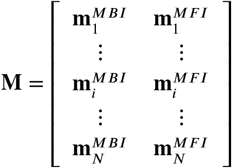

aganist which we perform a one-vs-rest classification of groups **y** with a sparse logistic regression classifier (implementing an *ℓ*_1_ penalty). Performance was measured using area the receiver operating characteristic curve (AUC) with a stratified-17-fold cross-validation procedure. A sparse model was chosen such that we could interrogate the coefficients for feature importance post-hoc. The number of cross-validation folds were chosen to be maximal with the constraint that each fold contained at least 2 classes in the validation partition. This analysis was done using the LogisticRegression class in the sklearn Python package (***Pedregosa et al., 2012***).

### Analysis of Reaction Times

The response variable in these analyses is the average reaction time difference between second-step choices made after rare and common trials: Δ_*RT*_ = 〈***RT*_*r*_〉** - 〈***RT*_*C*_**〉, where 〈·〉 denotes an average over trials of the subscripted type. Greater values of **Δ**_RT_ suggest greater reliance on MB control, since an unexpected outcome of a planned (MB) action likely induces some surprise, and thus slowing of the action at the second step. We fit a linear regression models of **Δ_RT_** against MBI and MFI between groups, and evaluated relationship magnitude and significance using the induced linear correlation coefficients.

### Reinforcement Learning Model

We selected the model architecture herein based on the results of our theory-free analysis, where the parameters of the RL model would offer further clarification.

Reinforcement learning models such as these can be split into two-components: a learning model, which describes the evolution of latent states of the decision-making system (i.e. the subject’s internal value function), and an observation model which describes how values represented in the learning model are transformed into observable behaviours (***Daw, 2011***). The learning model here assumes existence of two parallel systems, each of which learn representations of the expected value **a**^T^**s**_t_ of an action a taken when the agent is in a given state **s**_t_. The first system is a model-free learner, which is modeled according to the SARSA(1) algorithm [34]. Letting Q^π^ ∈ ℝ^2×3^ denote the model-free state-action value matrix, and superscripting states and action according to the step **s**^(j)^, **a**^(j)^ (i.e. for steps ***j*** = 1 or 2), we have the following updates.

First, the reward prediction error is calculated:

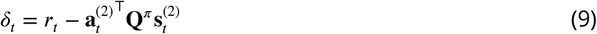

Second, the first model-free update based on Step 1 state and action:

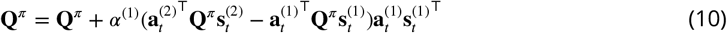

Third, the second step model-free update:

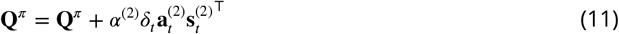

Finally, the second model-free update for the first step state and action:

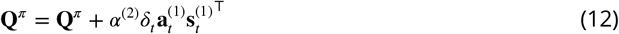

Here, states and actions are one-hot vectors of dimension 3 x 1 and 2×1, respectively. The **δ**_t_ is the reward prediction error at trial ***t***. The second element of the learning model is a MB learner, whose MB action values for step 1, Q^ω^ ∈ ℝ^2×1^, are computed at each trial using explicit representation of the state transition matrix and the Bellman equations [28,38].

The observation model dictates that the probability of a given action, ***p***_**a**_, at the first-step (dropping some superscripts for notational parsimony) is

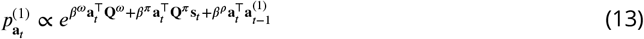

while at the second step it is

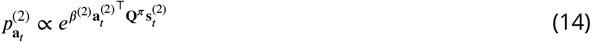

The inner product 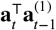 in Equation ***Equation 13*** is a binary indicator function which takes value 1 if the action at the first step of the present trial is identical to that taken at the first step on the previous trial. This reflects the subject’s tendency to perseverate (i.e. to make decisions simply based on repetition, irrespective of value). The symbols ***β*** = {***β***^*ω*^,***β***^*π*^,***β***^*ρ*^} represent weights on the model-based, model-free, and perseveration functions, respectively. At step-2, there is no model-based control (since this is the terminal step of the trial), and thus the inverse softmax temperature ***β***^(2)^ represents “choice consistency,” where lower values denote a tendency toward more stochastic decisions.

Each subject was thus characterized by a set of parameters 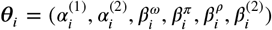 which we modeled as subject-level random-variables, ***θ***_*i*_ ~ ***P***(θ_*i*_|**ø**), with group-level priors **ø** = (***μ,σ***). Here ***μ*** ~ Cauchy(0,5), and ***σ*** ~ Cauchy(0,5) represent the group-level prior means over parameters = 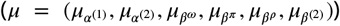 and the group level variance over each parameter 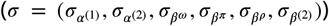 While the ***β*** parameters were all assumed to be normally distributed—and thus unconstrained in range—at the subject level (e.g. 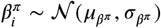 the learning rates ***α*** were constrained to lie on the (0,1) interval. Specifically, we sampled unconstrained values for the learning rates from a normal distribution, e.g. 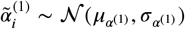 and subsequently used the inverse probit transform, yielding the constrained learning rate 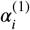 The same procedure was done for the second step learning rate.

We also introduced parameters to model the group level differences, as in ***Sharp et al. (2015)***. If we define [***BED***_i_] as a binary indicator function that takes value 1 if subject ***i*** is in the BED group and 0 if she is not, then we can define 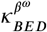,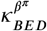,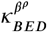 and 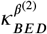 as the group level effect of BED on the model-based, model-free, and perseveration weights, ***β***^*ω*^, ***β***^*π*^,and ***β***^*σ*^ respectively, as well as the step-2 inverse softmax temperature ***β***^(2)^. We also define similar parameters for the BN group by simply replacing the subscript. The resulting hierarchical random-effects model can be summarized as follows

Group-Level:

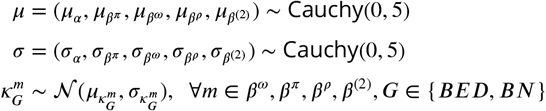

Subject-level:

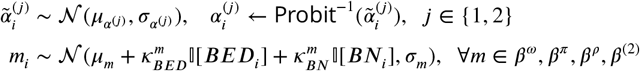

We performed inference using Markov chain Monte-Carlo in the Stan probabilistic programming language (***Carpenter et al., 2017***). The quality of parameter estimates was ascertained by visual inspection of the Markov-Chain convergence (via trace plots) as well as evaluation of the ***r̂*** statistic (***Gelman and Rubin, 1992***) (which was below 1.02 for all estimates in this model). Group-level differences were classified as such by determination of whether the respective 95% highest posterior density intervals for the group effect parameters crossed the origin, analogous to the frequentist interpretation of 95% confidence intervals.

### Principal Components Analysis

We sought to visualize the distribution of subject groups on a space defined by parameter estimates from both theory-free and RL analyses, as well as the reaction time analysis. We then projected the original data onto a 2-dimensional spaces defined by the first 3 principal components and evaluated the relative alignment of each model’s parameter estimates by visualization (there were insufficient data to perform confirmatory factor analysis). This was done using principal components analysis in the `factoextra package` (https://www.sthda.com/english/rpkgs/factoextra) for the R statistical programming language.

### Implementation Details

Analyses were performed on a Linux server with a 16-core AMD 1950X central processing unit. Theory-free analyses were conducted using the lme4 package (***Bates et al., 2015***). Reinforcement learning models were built and fit using the Stan programming language (***Carpenter et al., 2017***) and called within its Python interface. Visualizations were created using the R statistical programming language.

## Acknowledgments

The authors wish to acknowledge funding provided by the Nova Scotia Health Authority Research Fund and the Dalhousie Department of Psychiatry. The authors also acknowledge helpful feedback from Alexander Rudiuk and Dr. Valerie Voon, as well as Drs. Nathaniel Daw, Nuria Donamayor, and Valerie Voon for providing code for the task implementation.

## Appendix 1 Supplementary Figures

**Appendix 1 Figure 1.**
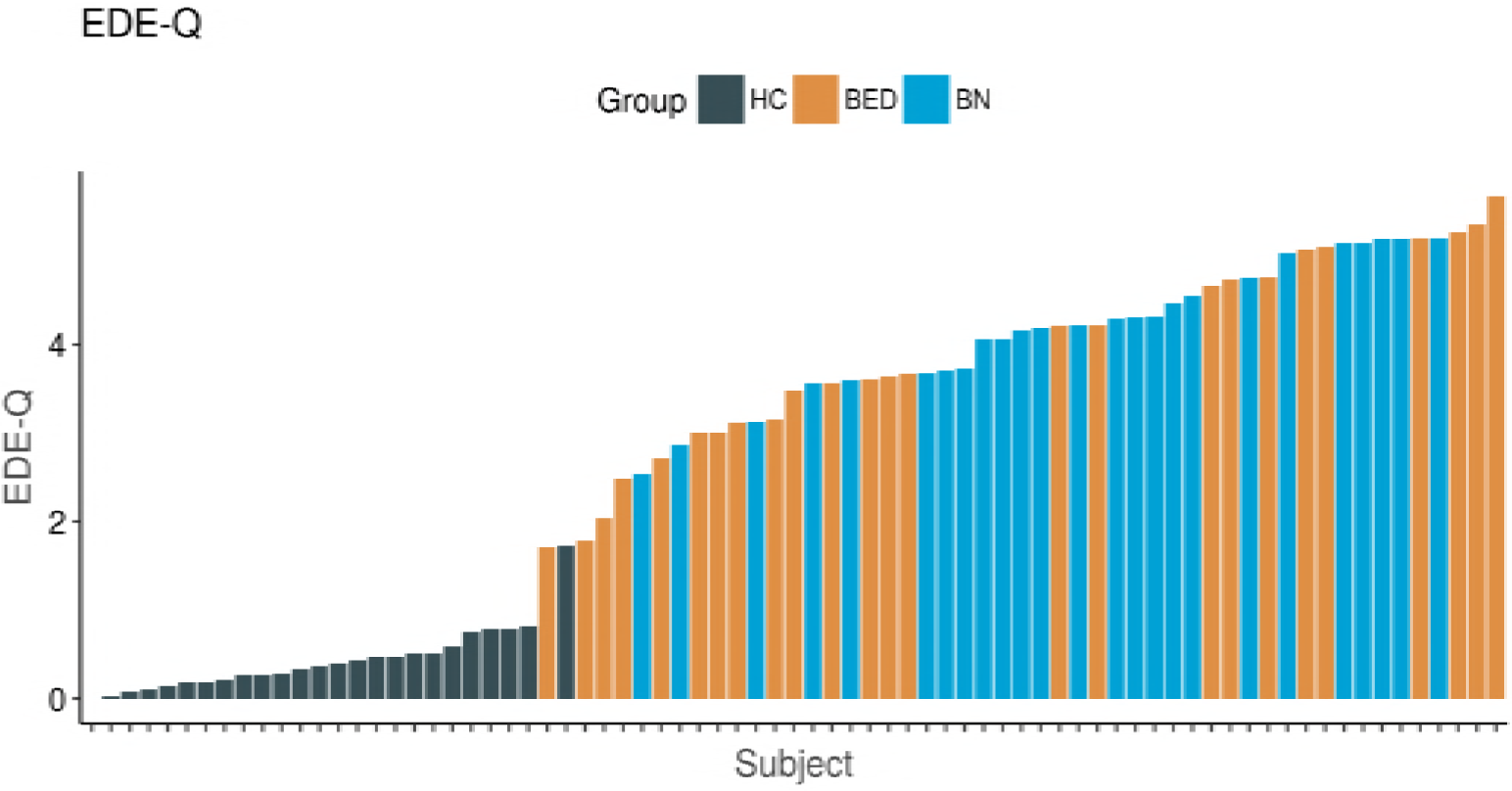
Distribution of EDE-Q scores across groups. Abbreviations: eating disorders examination questionnaire (EDE-Q), healthy control (HC), binge eating disorder (BED), bulimia nervosa (BN). Subjects are represented along the x-axis, and ordered by increasing EDE-Q score. Note the relatively sharp demarcation between healthy controls and the clinical groups. Among the disordered eating groups, there is a relatively similar distribution across EDE-Q scores.

**Appendix 1 Figure 2.**
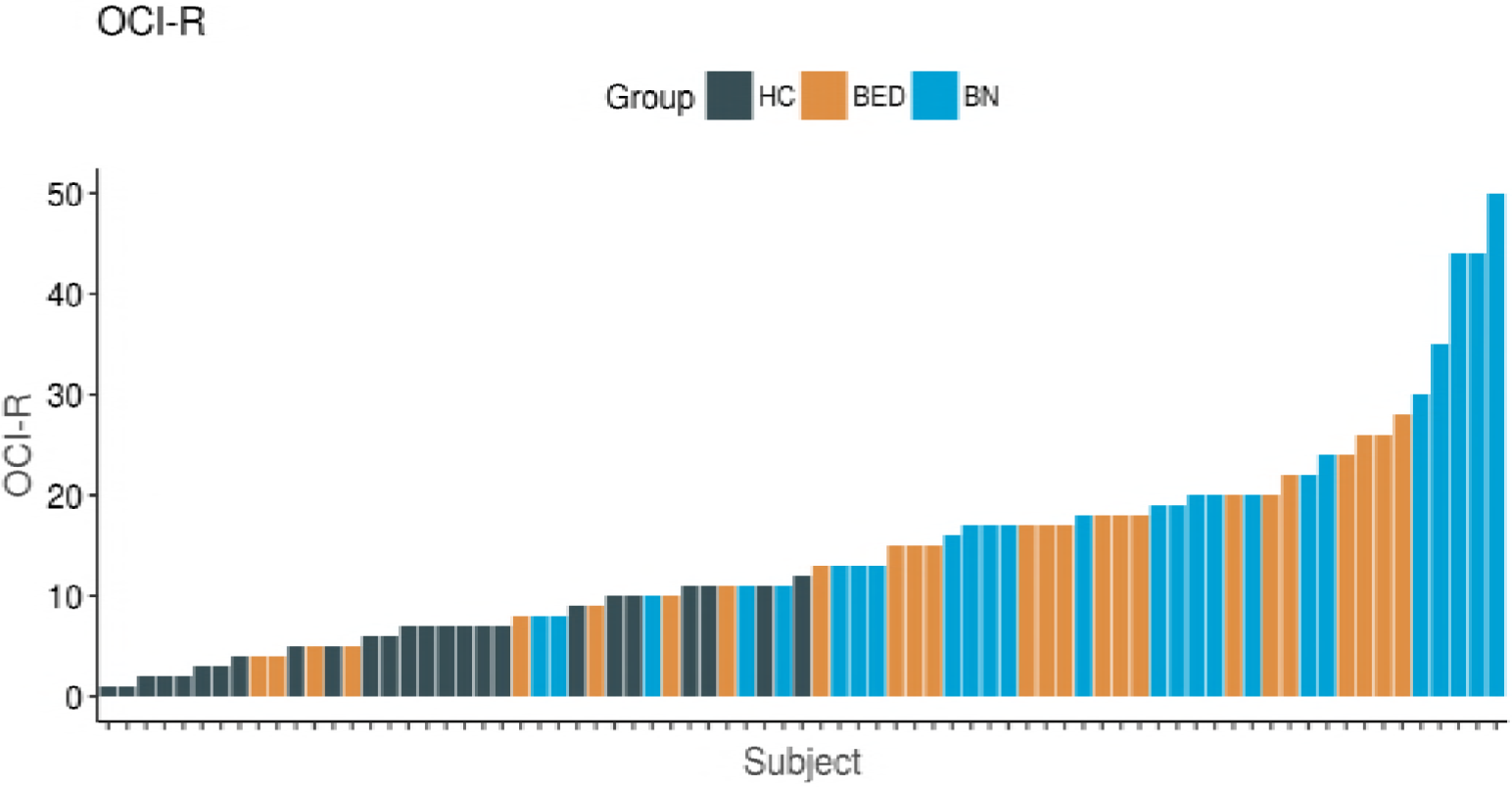
Distribution of OCI-R scores across groups. Abbreviations: obsessive-compulsive inventory-revised (OCI-R), healthy control (HC), binge eating disorder (BED), bulimia nervosa (BN). Subjects are represented along the x-axis, and ordered by increasing OCI-R score. Pathological binge eating groups demonstrated higher scores on the OCI-R relative to healthy controls, with the most extreme scores found in a subset of BN subjects.

**Appendix 1 Figure 3.**
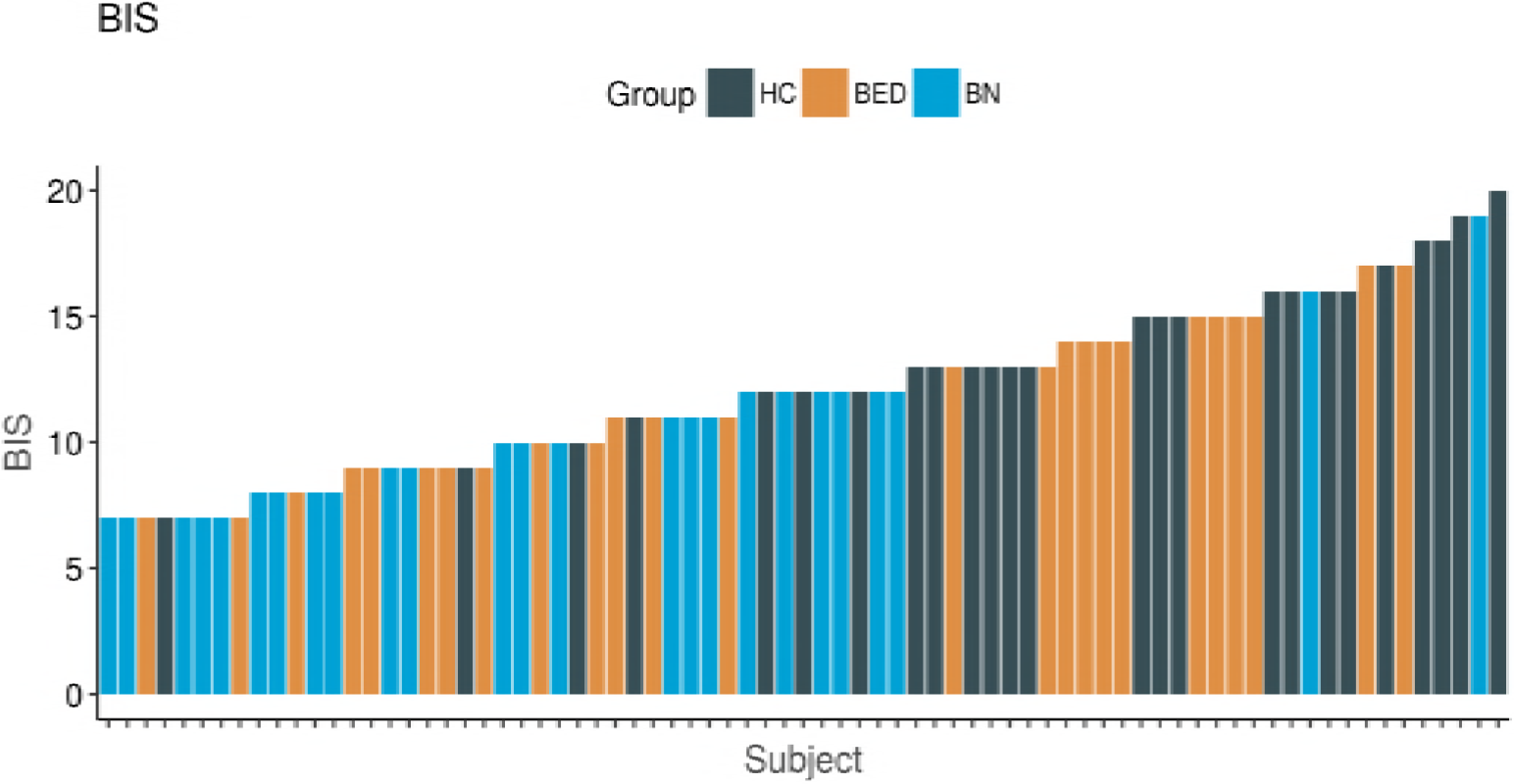
Distribution of BIS scores across groups. Abbreviations: behavioural inhibition scale (BIS), healthy control (HC), binge eating disorder (BED), bulimia nervosa (BN). Subjects are represented along the x-axis, and ordered by increasing BIS score. There is relatively less sharp demarcation of the healthy controls and pathological binge eating group along this measure, although healthy controls had statistically significantly higher scores in general.

**Appendix 1 Figure 4.**
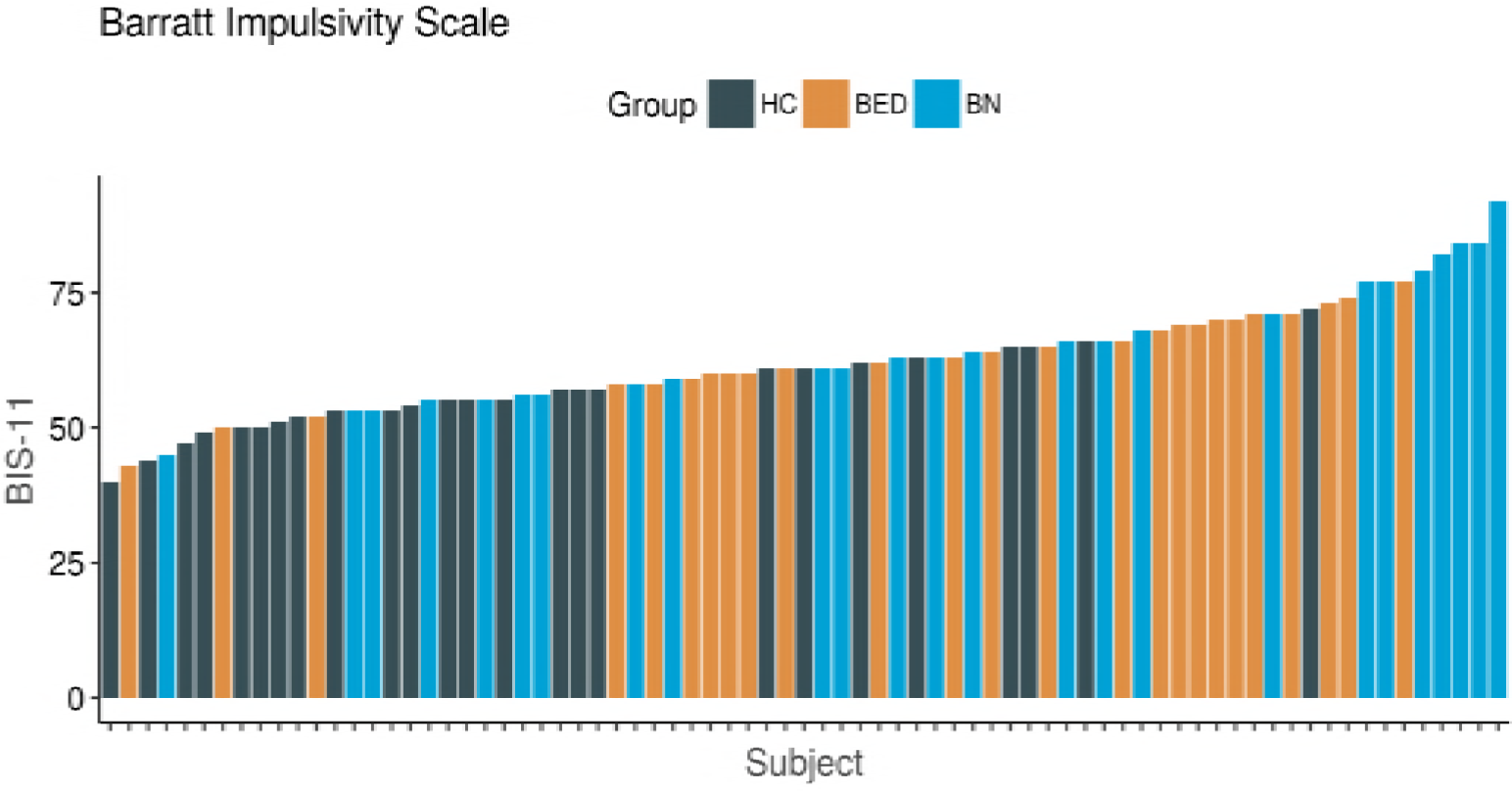
Distribution of BIS-11 scores across groups. Abbreviations: Barratt impulsivity scale (BIS-11), healthy control (HC), binge eating disorder (BED), bulimia nervosa (BN). Subjects are represented along the x-axis, and ordered by increasing BIS-11 score. There is relatively less sharp demarcation of the healthy controls and pathological binge eating group along this measure, although pathological binge eating groups had statistically significantly higher scores in general relative to healthy controls.

**Appendix 1 Figure 5.**
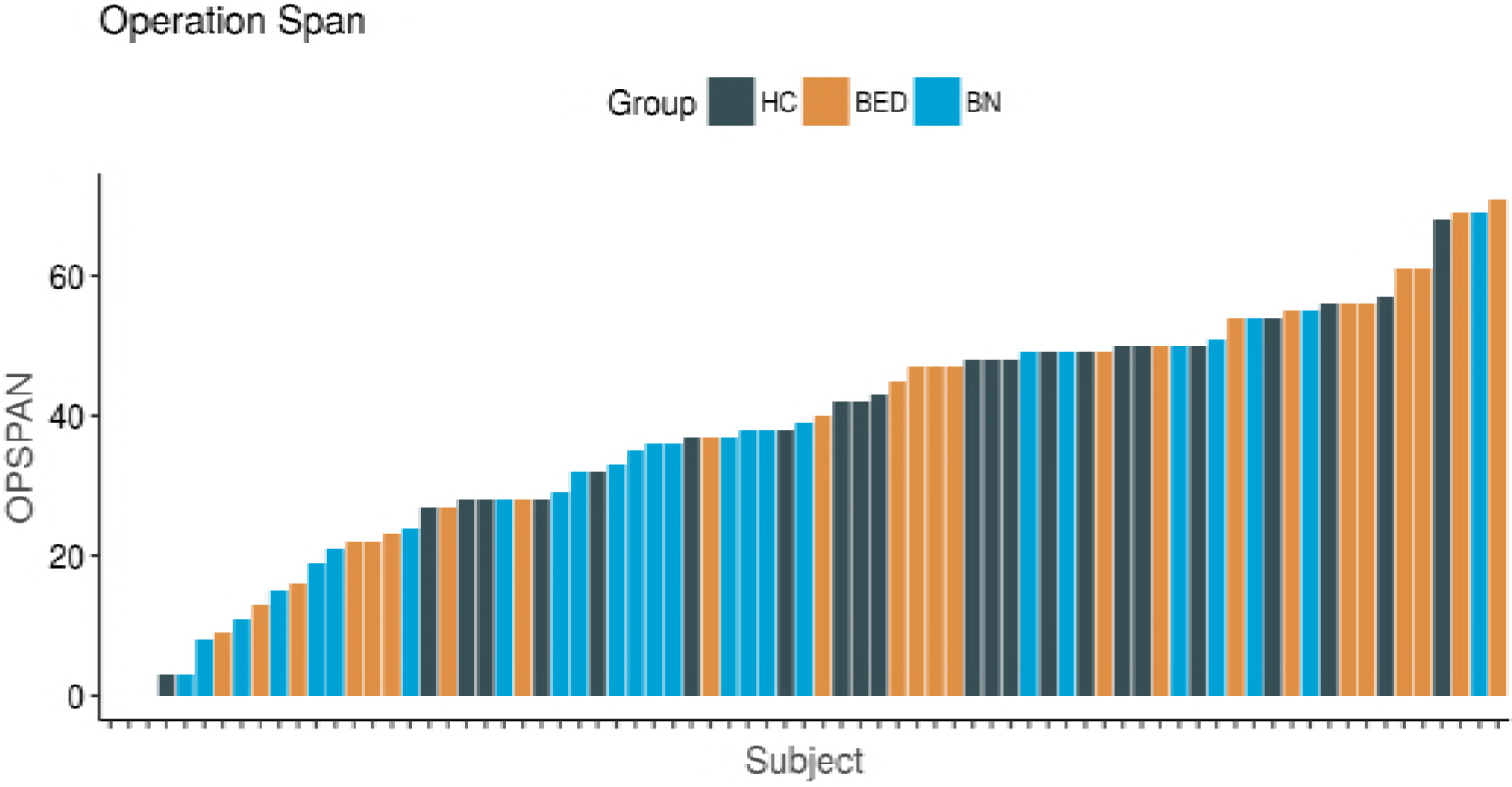
Distribution of OPSPAN scores across groups. Abbreviations: operation span (OPSPAN), healthy control (HC), binge eating disorder (BED), bulimia nervosa (BN). Subjects are represented along the x-axis, and ordered by increasing OPSPAN score. There was no significant difference between groups in terms of working memory.

